# Citrullination of a phage displayed human peptidome library reveals the fine specificities of rheumatoid arthritis-associated autoantibodies

**DOI:** 10.1101/2021.04.22.441021

**Authors:** Gabriel D. Román-Meléndez, Daniel R. Monaco, Janelle M. Montagne, Rachel S. Quizon, Maximilian F. Konig, Mekbib Astatke, Erika Darrah, H. Benjamin Larman

## Abstract

Post-translational modifications (PTMs) on proteins can be targeted by antibodies associated with autoimmunity. Despite a growing appreciation for their intrinsic role in disease, there is a lack of highly multiplexed serological assays to characterize the fine specificities of PTM-directed autoantibodies. In this study, we used the programmable phage display technology, Phage ImmunoPrecipitation Sequencing (PhIP-Seq), to profile rheumatoid arthritis (RA) associated anti-citrullinated protein antibody (ACPA) reactivities. Using both an unmodified and peptidylarginine deiminases (PAD)-modified phage display library consisting of ~250,000 overlapping 90 amino acid peptide tiles spanning the human proteome, PTM PhIP-Seq robustly identifies antibodies to citrulline-dependent epitopes. PTM PhIP-Seq was used to quantify key differences among RA patients, including PAD isoform specific ACPA profiles, and thus represents a powerful tool for proteome-scale antibody-binding analyses.

## Introduction

Phage ImmunoPrecipitation Sequencing (PhIP-Seq) is a programmable phage display technology that enables unbiased analysis of antibody binding specificities.^1,2,3^ An oligonucleotide library encoding the complete human proteome as ~250,000 overlapping 90 amino acid peptides was cloned into the mid-copy T7 phage display vector (~5-15 copies of displayed peptide per phage particle) and is immunoprecipitated by serum antibodies for analysis by high throughput DNA sequencing. This enables serum antibody profiling against hundreds of thousands of peptide epitopes at low cost.^2,4^ However, since the phage library is produced in *E. coli*, the displayed peptides lack post-translational modifications (PTMs) relevant to human disease.^5,6^ Here, we enzymatically modify the phage-displayed human peptidome, such that PTM-dependent autoantibody reactivities can be precisely assessed by quantitative comparison to the unmodified library.

Citrullination is the post-translational conversion of arginine in proteins to citrulline, which in humans is catalyzed by the calcium-dependent peptidylarginine deiminase (PAD) family of enzymes.^7,8^ This modification can impart antigenicity to self-proteins, and in the case of rheumatoid arthritis (RA), results in the production of anti-citrullinated protein antibodies (ACPAs).^9^ These antibodies are believed to participate in disease pathogenesis by binding citrullinated proteins in the synovial joint.^10,11^ The discovery of autoantibodies to peptidylcitrulline epitopes in RA serum led to the development of a diagnostic Enzyme Linked Immunosorbent Assay (ELISA) using synthetic cyclic citrullinated peptides (CCP).^12^ Since this anti-CCP antibody test can achieve diagnostic sensitivity of ~80% and a specificity up to 98%, anti-CCP antibodies have become a routinely utilized biomarker for RA.^13,14^

Of the five known PAD isoforms, PAD2 and PAD4 make up the majority of expressed PAD in inflamed synovial tissue.^15^ Studies using recombinantly expressed enzymes suggest that PAD2 and PAD4 can generate distinct, yet overlapping sets of epitopes that can be targeted by ACPAs.^16,17,18^ ACPAs that recognize these substrates are known to cross-react, making it difficult to distinguish the relative contributions of the different PAD isoforms to the expression of ACPAs.^19,20,21^ While aberrant citrullination is best known for its role in RA pathology, it is also implicated in other disease processes, including multiple sclerosis, Alzheimer’s disease, and cancer.^22,23^ Therefore, unbiased identification of antibody reactivities to citrulline epitopes is of broad interest.

## Results

### Citrullination of the T7 phage displayed human peptidome library

In a typical PhIP-Seq experiment, the phage display library is mixed with 0.2 μl of serum or plasma and immunoprecipitated (IPed) using magnetic beads coated with protein A and protein G, which together capture all IgG subclasses. The DNA sequences from the immunoprecipitated phage clones are then amplified by PCR and the abundance of each clone is quantified by high throughput DNA sequencing. To detect citrullinated peptide autoreactivities, we first developed a method to enzymatically citrullinate the human 90-mer phage library^24^ *in vitro* using recombinant human PAD2 and PAD4 enzymes (Methods). Sequencing read count data from PhIP-Seq with a citrullinated library can be directly compared against count data from PhIP-Seq with an unmodified library. **Fig. 1** illustrates how this type of analysis is able to identify citrulline-dependent autoantibody binding specificities.

**Figure 1.**
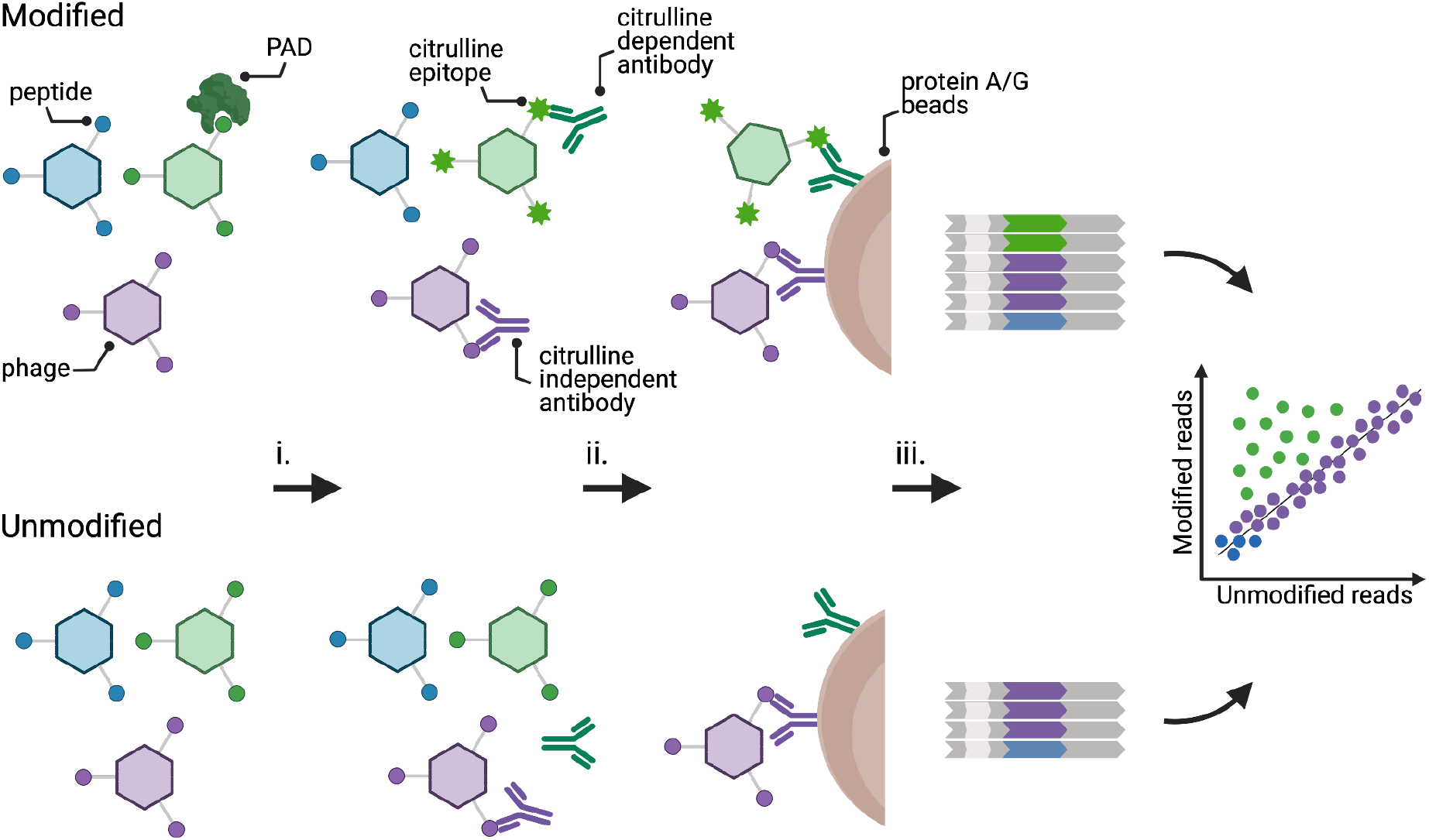
Experimental design of PAD PhIP-seq. (i) PAD-modified (top) and unmodified (bottom) phage libraries are mixed with patient serum. (ii) The resulting phage-antibody immunocomplexes are then precipitated on Protein A/G beads and unbound phage washed away. (iii) Peptide inserts of the immunoprecipitated clones are PCR amplified and analyzed by high throughput DNA sequencing. Abundant clones along the diagonal (purple) represent reactive peptides that do not depend upon modification. Off-diagonal peptides (green) are immunoprecipitated only after being post-translationally modified. Low abundant clones and non-specific binding cluster at the origin (blue).

### Phage display library modification and quality assessment

We reasoned it would be important to establish a rigorous and generalizable quality control pipeline for assessing the integrity of PTM phage libraries. We first confirmed that buffer exchange using standard dialysis into two common buffers (PBS and TBS) did not impact phage viability (**Fig. S1a**). Second, we assessed phage viability after enzymatic modification. The 90-mer library, a control clone that does not display a peptide, and a second control clone that displays a peptide that is not a PAD substrate, were all found to retain their infectivity after treatment with PAD enzyme (**Fig. S1b**). We also confirmed by western blot that the buffer and conditions used to citrullinate the phage display library were compatible with robust PAD activity (**Fig. S1c**).

We next sought to confirm that PAD does not create reactive citrulline epitopes on the phage coat proteins. If it did, the entire library might be non-specifically immunoprecipitated by ACPAs. To this end, we assessed the binding of anti-CCP+ versus anti-CCP- sera to a phage clone without a displayed peptide, which was either subjected to PAD4 modification or not. Anti-CCP+ serum did not precipitate this modified phage clone (**Fig. S1d**). Lastly, we confirmed that treatment with PAD enzyme does not interfere with PCR amplification of the library inserts (**Fig. S1e**). In summary, the phage library can be successfully citrullinated *in vitro* using recombinant PAD enzyme without loss of viability and without the formation of phage capsid citrulline epitopes, while remaining compatible with downstream amplification for sequencing. A flow chart illustrating this quality control pipeline, which is readily generalizable to other PTM PhIP-Seq studies, is shown in **Fig. S2**.

We next analyzed negative control (“mock”) IPs that omitted antibody input in order to characterize the background binding of each modified and unmodified library peptide to the protein A/G-coated magnetic beads. The mock IPed libraries were sequenced to a depth of ~20-fold. The mock IPs of the PAD2- and PAD4-modified libraries exhibited an essentially indistinguishable background binding profile compared to the unmodified library (**Fig. S3**). Conveniently, pairwise comparison of modified and unmodified libraries is therefore not confounded by bias in background binding to the beads, simplifying downstream analyses.

### PAD PhIP-Seq robustly identifies antibodies to citrulline modified epitopes

We next sought to detect binding of PAD-dependent peptide epitopes by ACPAs from anti-CCP+ RA sera. Unmodified, PAD2-modified, and PAD4-modified libraries were separately immunoprecipitated with RA51, an anti-CCP+ serum sample. The PhIP-Seq read count data of RA51 was then subjected to a pairwise analysis pipeline^25^ to identify PAD-dependent antibody reactivities. Comparison of the unmodified versus PAD-modified libraries identified 294 PAD2- and 306 PAD4-dependent reactivities (**Fig. 2a-b, Table S1**). In total, 427 unique peptides were found to exhibit PAD-dependent reactivity; 121 modified peptide reactivities were specific to PAD2, whereas 133 were specific to PAD4. The remaining 173 modified peptide reactivities were shared among both enzymes (**Fig. 2c**).

**Figure 2.**
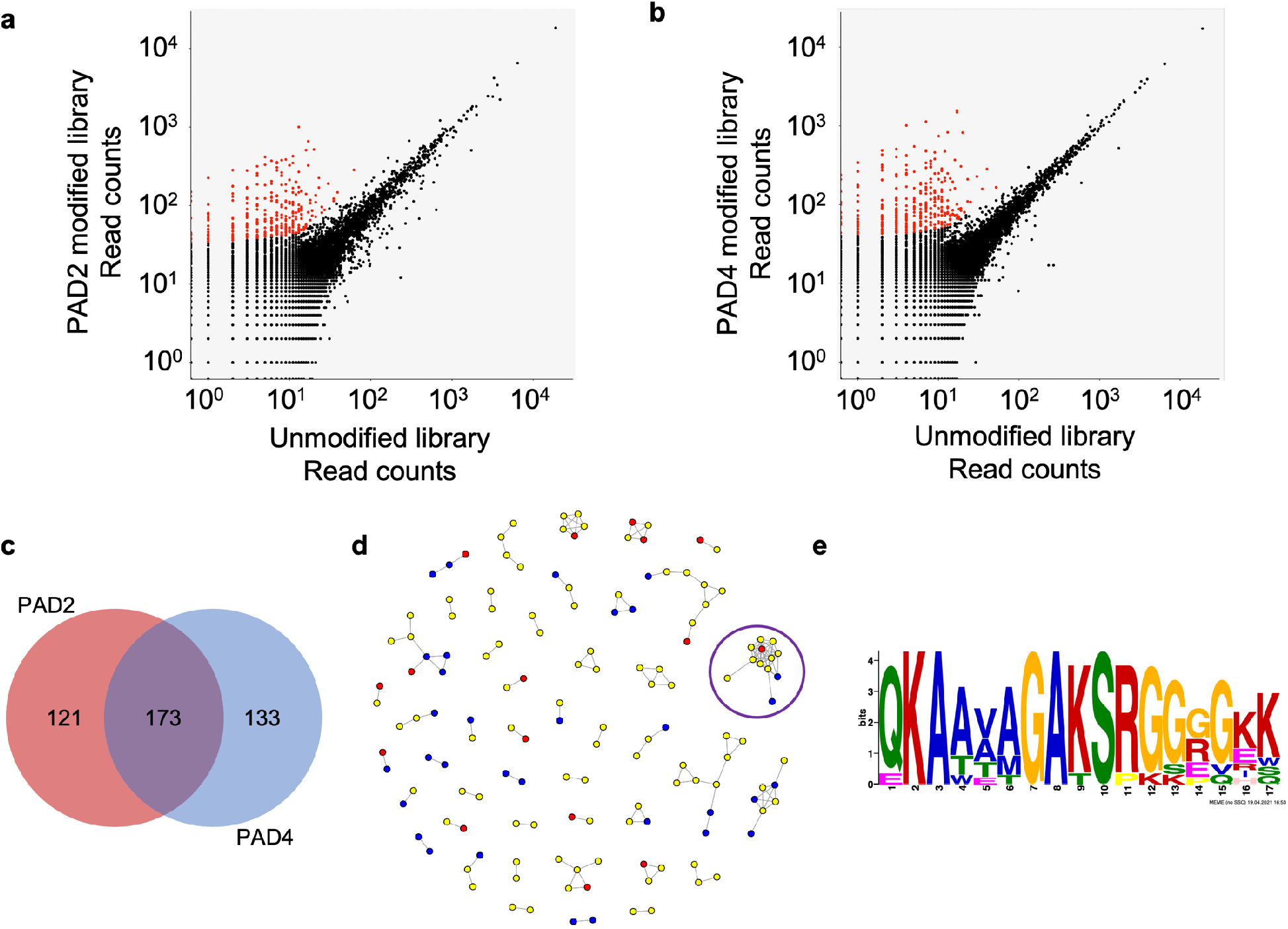
Profiling PAD-dependent antibody reactivity. **a** Scatter plot comparison of PAD PhIP-Seq data from separate IPs of the unmodified and the PAD2-citrullinated and **b** PAD4-citrullinated library using serum from RA51, an anti-CCP+ RA patient. Citrulline-dependent peptide reactivities are colored red. **c** Pie chart demonstrating the distribution of citrulline-dependent epitopes identified from PAD PhIP-Seq of RA51. **d** Network graph of all reactive peptides in **a** and **b**. Nodes represent reactive peptides from the PAD2 library (red), PAD4 library (blue), or both PAD2 and PAD4 libraries (yellow). Nodes are linked if the peptides share sequence homology. The purple circle indicates the largest cluster. **e** A multiple sequence alignment logo generated from the largest cluster in **d**.

All RA51 serum reactive peptides were searched for shared sequences using the shared epitope detection algorithm epitopefindr.^26^ The results are visualized as a network graph in which peptides are represented as nodes and a potentially shared epitope as edges (**Fig. 2d**). Peptides in the largest cluster were then analyzed using the motif discovery software MEME,^27^ which revealed a conserved sequence modified by both PAD2 and PAD4. The motif features -RGGGGK- and likely contains the citrullinated arginine (**Fig. 2e**).^27^ Crystallographic data suggests that PAD4 recognizes five successive residues with a ØXRXX consensus where Ø represents amino acids with small side-chains and X denotes any amino acid.^28^ This is in agreement with an *in silico* -RXXXXK-PAD_2_ model and a -RGXXXX-PAD_4_ model (with a strong preference for glycine at the +1, +2, and +3 positions),^29^ as well as the experimentally determined PAD4 citrullination sequences -RG/RGG- of FET proteins^30^ and -SGRGK- of histone H2A.^31^ PTM PhIP-Seq can thus be used to characterize the substrate specificity of enzymes imparting post-translational modifications.

### Longitudinal analysis of ACPA fine specificities using PAD PhIP-Seq

ACPA are expressed at all stages of RA and can precede the onset of clinical disease by over a decade.^32^ Clinically, ACPAs are routinely used to aid in the diagnosis of RA and provide prognostic information about disease severity and phenotype (e.g., risk of erosive disease and interstitial lung disease).^33^ Moreover, changes in anti-CCP levels correlate with treatment responses.^34^ By monitoring ACPA fine specificities, however, it may be possible to increase the sensitivity and specificity of this diagnostic modality. As a proof of concept, we longitudinally profiled serum from an anti-CCP+ RA patient with high disease activity. PAD4 PhIP-Seq was employed to probe the individual’s ACPA repertoire at seven timepoints, both pre- and post-treatment with the B cell depletion therapy, rituximab (**Figs. 3, S4**). Reactivity profiles obtained prior to treatment harbored consistent levels of PAD4-dependent reactivity, directed at ~100 citrullinated peptides comprising ~80 distinct protein targets. Subsequent to B cell depletion, the number of PAD4-dependent reactivities declined at a rate of approximately 9 peptides per week for 9 weeks (R^2^=0.8 during this time period). This fall in citrulline-dependent antibody levels and diversity was followed by clinical improvement of synovitis as measured by Clinical Disease Activity Index (36 on Day 1 and 10.5 a month after the first rituximab treatment). These data indicate that PAD PhIP-Seq can be used to characterize the temporal evolution of citrulline-dependent autoantibody fine specificities in response to treatment.

**Figure 3.**
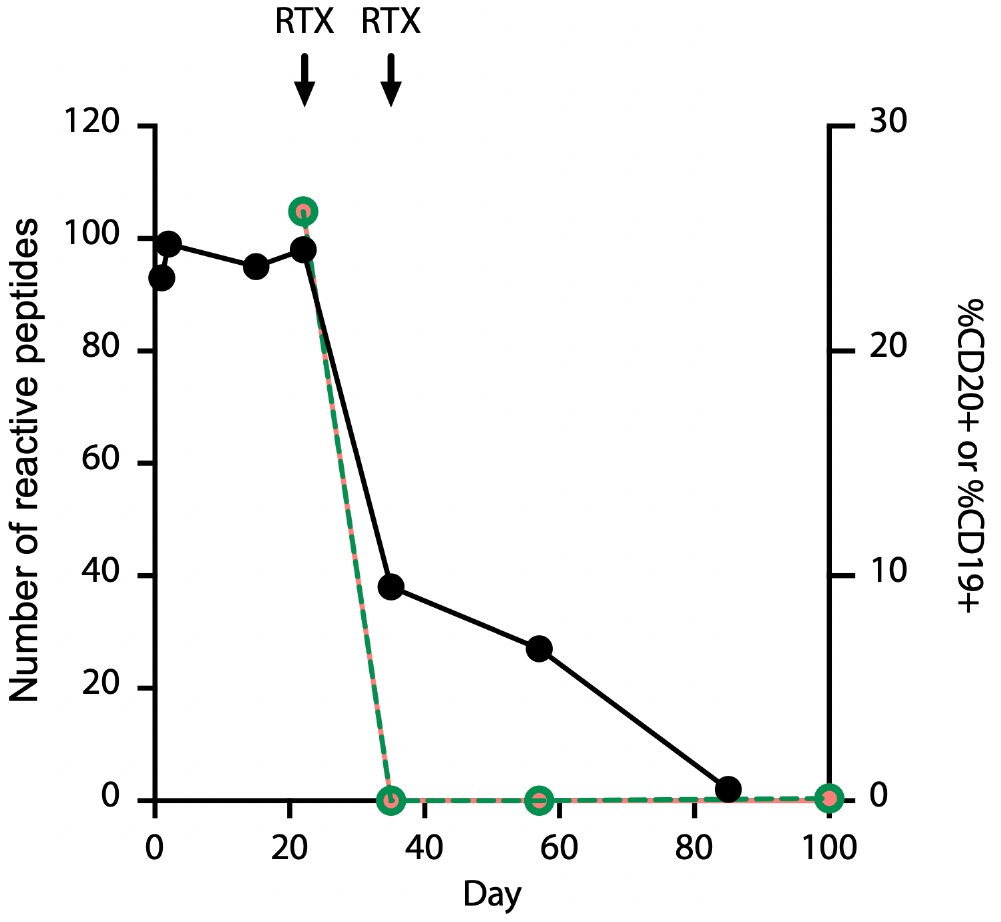
Longitudinal monitoring of ACPA reactivity in a patient with RA using PAD4 PhIP-seq. The total number of immunoprecipitated citrullinated peptides (black) declines following B cell depletion with rituximab (arrows) starting on Day 22, at which point a steady decline in citrulline-dependent antibody reactivity is observed at a rate of y=−1.3x+106.9 (R^2^=0.8) over the course of 9 weeks. The percentages of CD19+ (magenta) and CD20+ (green) cells of total lymphocytes in peripheral blood are shown on the secondary axis. Immunoprecipitated peptides on days 0-22 are reflective of baseline treatment with methotrexate and glucocorticoids.

### PAD PhIP-Seq positivity is concordant with clinical CCP test results

We next asked how well the PAD PhIP-Seq assay correlates with clinical anti-CCP serostatus. Unmodified, PAD2-modified and PAD4-modified libraries were separately immunoprecipitated using sera from a cohort of 30 RA patients and 10 demographically matched healthy controls. Upon sample unblinding, 20 of 21 anti-CCP+ patients were found to exhibit significant reactivity to a large set of PAD2 and/or PAD4-dependent peptide epitopes (**Fig. 4a-b, Table S1**). Using a cutoff of ≥15 PAD-dependent reactive peptides, PAD PhIP-Seq positivity was highly concordant with the clinical antigen-agnostic CCP assay; only one anti-CCP+ serum recognized neither PAD2 nor PAD4 PhIP-Seq peptides, and one additional anti-CCP+ serum recognized PAD4 PhIP-Seq peptides, but did not recognize PAD2 PhIP-Seq peptides above threshold (**Table S1**). It is important to note that the observed discordance may be due to CCP testing and PAD PhIP-Seq analysis not always being performed on the same sample timepoints. None of the sera from the 10 healthy controls, nor that of a Sjogren’s Syndrome patient, reacted with PAD PhIP-Seq peptides above threshold (**Table S1**). In this cohort, the PAD PhIP-Seq assay performed with a sensitivity of 90-95% and a specificity of 100%, when compared to the standard-of-care clinical assay that globally captures ACPAs using artificial CCP peptides.

**Figure 4.**
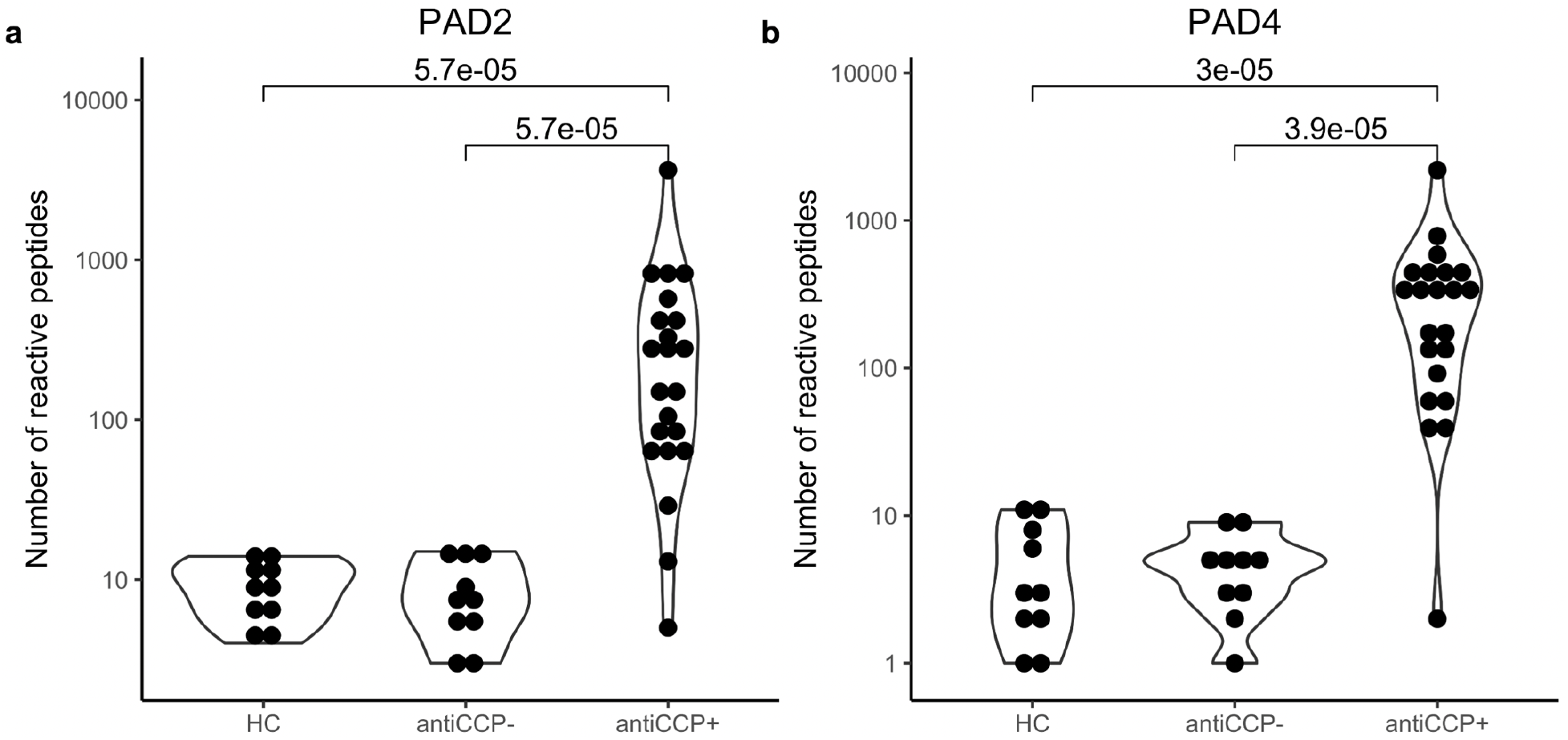
Cross sectional concordance of PAD PhIP-Seq with the clinical CCP assay. **a** The number of differentially reactive PAD2-dependent and **b** PAD4-dependent epitopes, per individual across the three groups: anti-CCP-RA, anti-CCP+ RA, and healthy controls (HC). A pseudo count of 1 was added to all individuals for plotting on a logarithmic scale.

### PAD PhIP-Seq reveals PAD2- and PAD4-dependent autoantibody reactivity signatures

Small molecule inhibitors of PAD enzymatic activity have emerged as promising candidates for the treatment RA.^35,36,18^ Pan-PAD inhibitors and those with distinct inhibitory profiles against PAD2 and PAD4 have been developed.^37^ It is therefore of great interest to determine whether subgroups of RA patients harbor ACPAs that exhibit a discernable preference for PAD2-versus PAD4-modified epitopes. Isoform-preferring antibody signatures, if they exist, may have utility in distinguishing subgroups of individuals more likely to benefit from one class of PAD inhibitors over another.

To identify potential PAD2- and PAD4-specific antibody signatures, we hierarchically clustered the anti-CCP+ patient subset based on their reactivity profiles (**Fig. 5a**). While we observe diverse reactivity to both PAD2- and PAD4-modified peptides, the clustering suggested the existence of distinct, patient-specific reactivity preferences among PAD2 versus PAD4 substrates. A total of 1,161 PAD-dependent peptides were reactive to at least three sera; 303 were unique to PAD2, 272 were unique to PAD4, and 293 peptides were reactive after modification by either enzyme (**Fig. 5b**). Among these dominant peptides are the known PAD substrates hornerin (HNNR)^38^, keratin (KRT4)^39^, and collagen (COL2A1)^40^. Other well-known substrates, however, including histone, vimentin, and fibrinogen, were not represented. The absence of these likely reflect a requirement for a higher level of conformational structure to serve as a PAD substrate and/or an antibody epitope, compared to the 90 amino acid peptides presented on the surface of the T7 phage.

**Figure 5.**
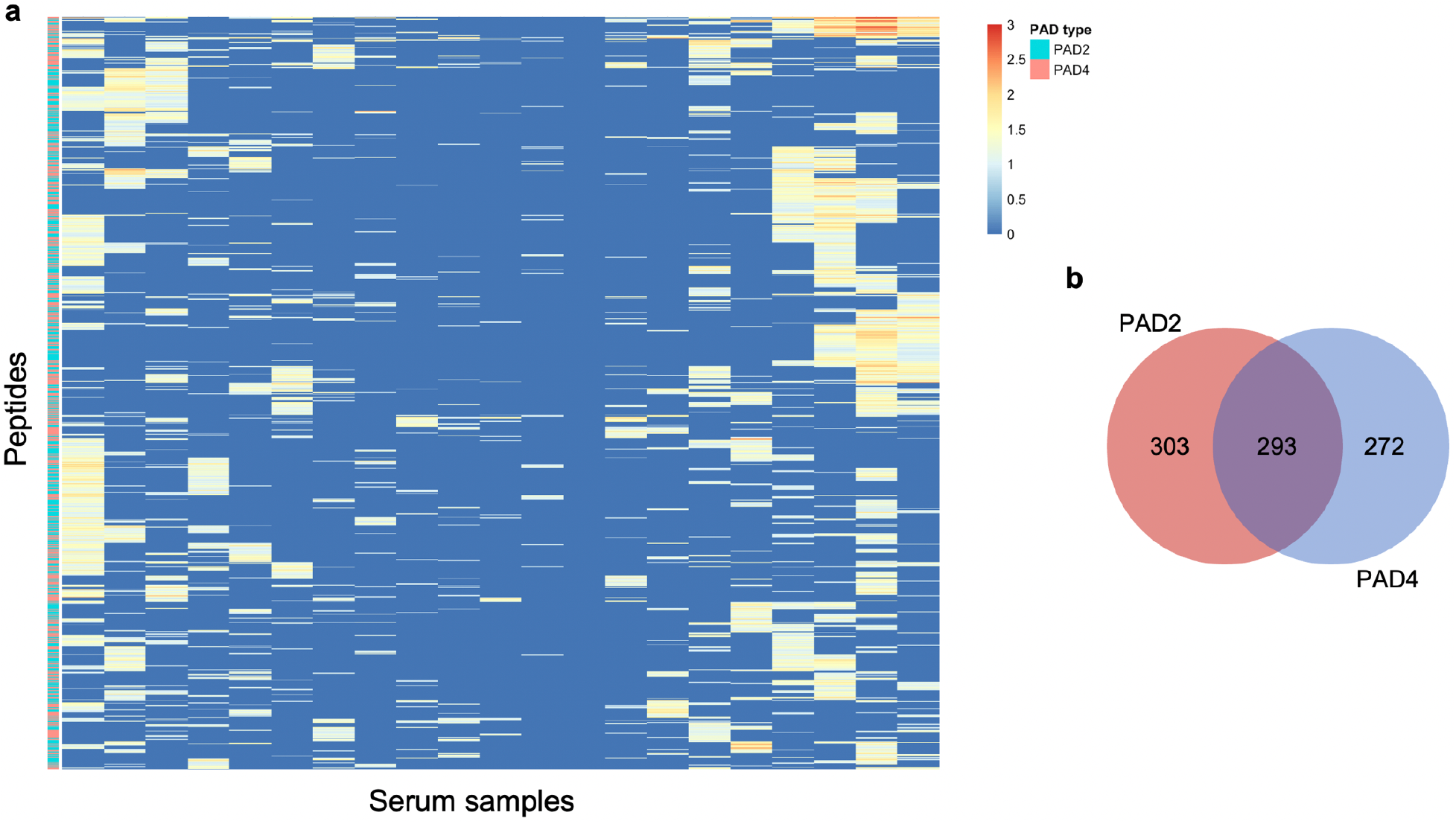
Reactivity profile of anti-CCP+ individuals. **a** Clustering of ACPA reactivity to citrulline-containing epitopes. Rows are peptides recognized by at least three anti-CCP+ sera (columns). **b** Distribution of the citrulline-dependent epitopes identified in **a**.

We observed that some of the dominant peptides appeared to be differentially reactive, depending on whether they were modified by PAD2 or PAD4. The number of anti-CCP+ individuals reactive to a given PAD modified peptide is shown in **Fig. 6a**, illustrating peptide dominance as a function of PAD2 versus PAD4 modification. Approximately 2.5% percent of all peptides were reactive in at least one anti-CCP+ individual. The most dominant PAD-dependent peptide reactivities were substrates that could be modified by either PAD2 or PAD4. Several peptides, however, were converted into citrullinated epitopes exclusively by one or the other enzyme: 65 by PAD2 and 50 by PAD4. We reasoned that these particular peptides could be used to probe the PAD specificity profile of ACPAs. A second potential application for these peptides would be as sensors for PAD2 or PAD4 enzymatic activity in a biological matrix.

**Figure 6.**
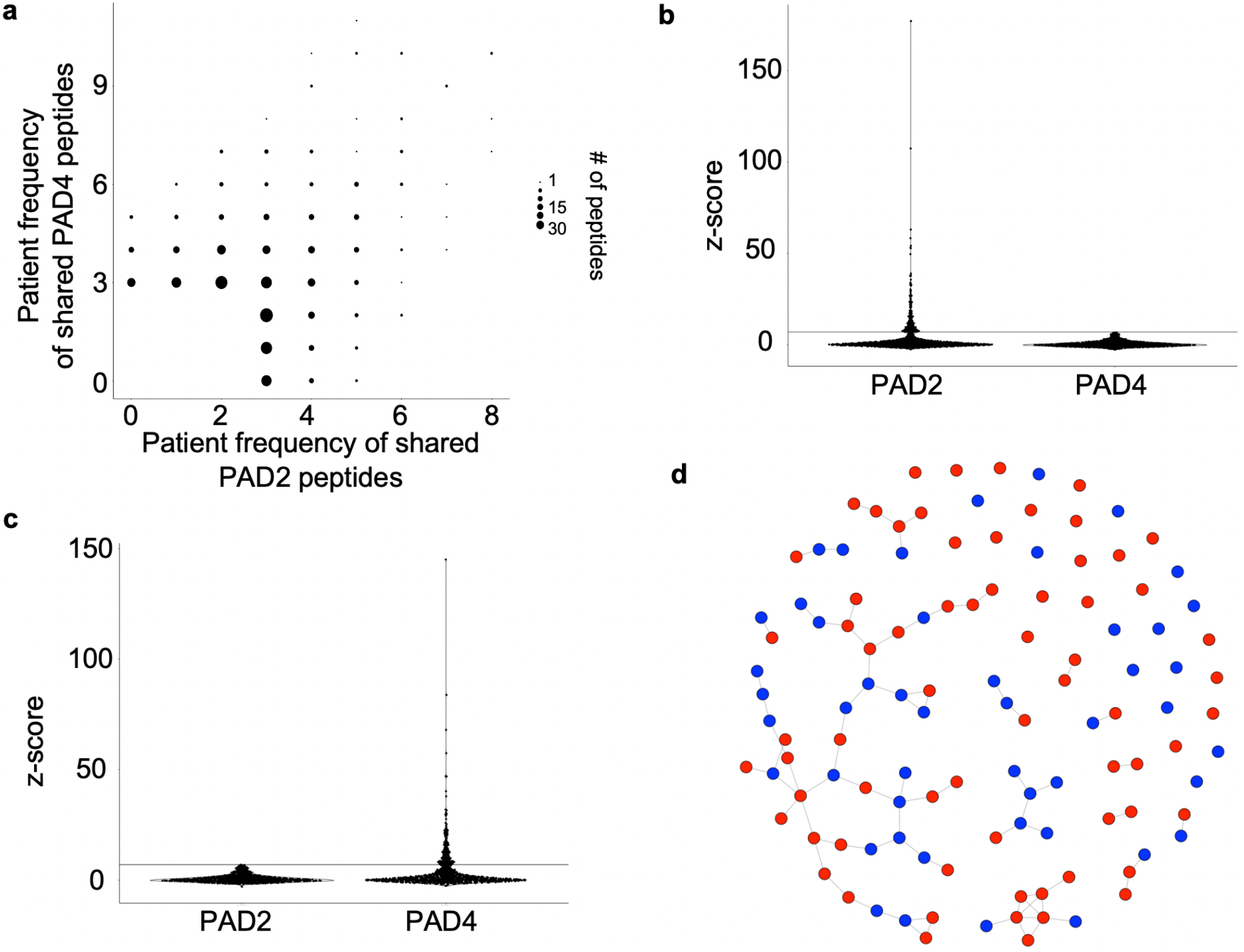
Identifying PAD2 and PAD4 reactivity signatures. **a** Number of anti-CCP+ individuals reactive to a PAD modified peptide. Data is shown for peptides (dots) reactive in at least 3 people. **b** Distribution of PAD2-specific and **c** PAD4-specific reactivities for all anti-CCP+ individuals. **d** Network graph of all PAD2 (red) and PAD4 (blue) specific reactive peptides. Nodes are peptides and are linked if they share sequence similarity.

The of PAD2- and PAD4-specific peptide reactivities across all the anti-CCP+ individuals are shown in **Fig. 6b-c.** Many peptides are strongly reactive after modification by one, but not the other, PAD enzyme (“mono-reactive”). A shared epitope network graph was used to characterize the degree of sequence homology among these 115 mono-reactive peptides. Approximately 50% of the mono-reactive peptides form nodal networks of >3 edges with the largest network consisting of 42 peptides (**Fig. 6d, Fig. S5**). A total of 32 mono-reactive peptides remains as singletons and do not align to any other mono-reactive peptide. A comparison of the number of PAD2-versus PAD4-specific reactivities detected in each anti-CCP+ individual is shown in **Fig. 7**. Overall preference for PAD2 versus PAD4 was quantified using an exact binomial test. A significant differential PAD2/4 “preference” was detected in some anti-CCP+ individuals (**Table S2**). These data support further investigation into specific PAD isoform-associated ACPA profiles and their potential relevance for prognostication and/or PAD-inhibitor companion diagnostic testing.

**Figure 7.**
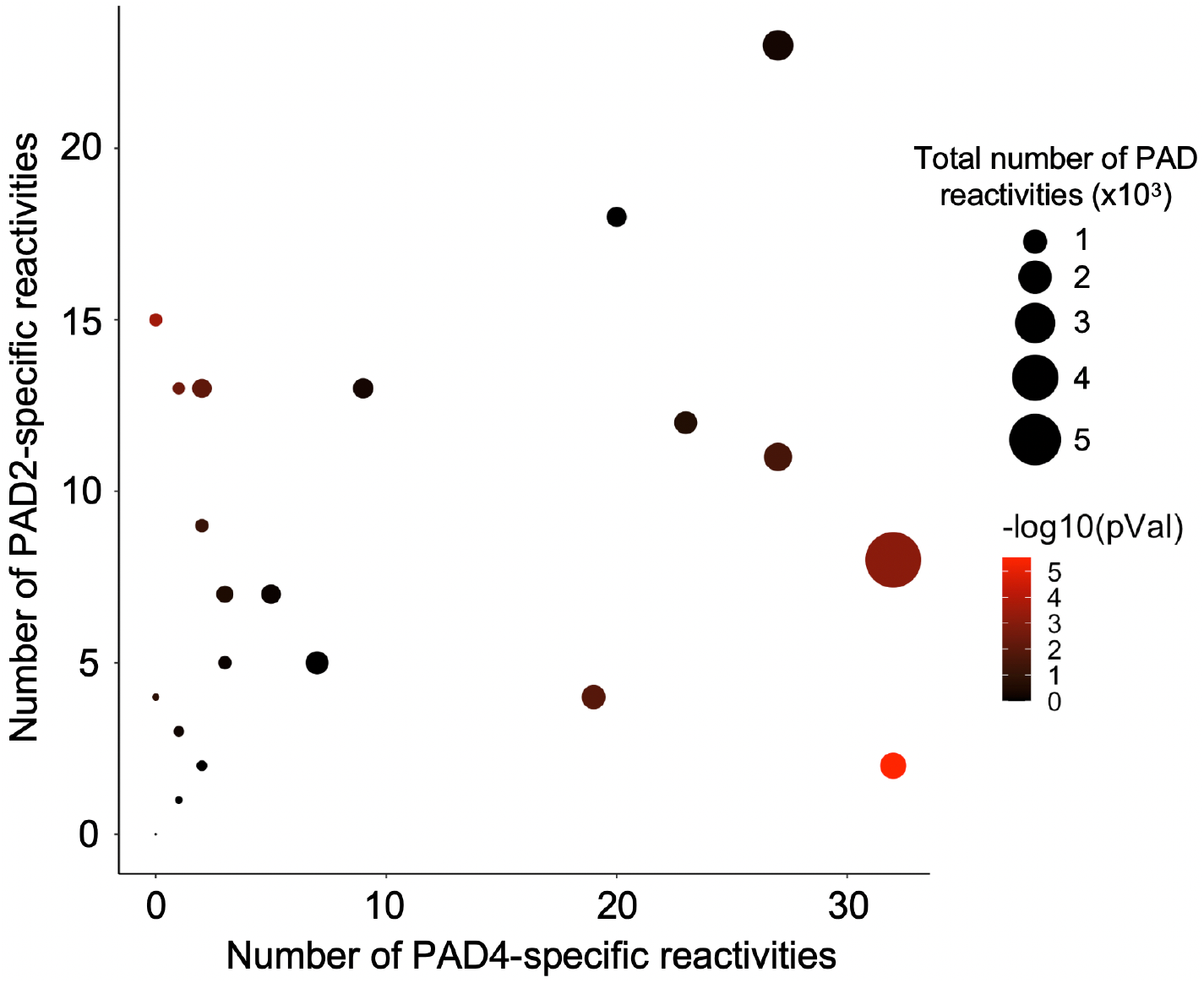
Number of PAD specific reactivities for all anti-CCP+ individuals. Dot size is proportional to the total number of PAD2 and PAD4 reactivities. Statistical significance was determined using an exact binomial test assuming a hypothesized null probability of 0.5 for independent peptides and is represented as a color gradient from black (not significant) to red (significant).

## Discussion

Serum antibody profiling can inform understanding of human immune processes, as well as suggest disease-specific antibody biomarkers. Despite advances in highly multiplexed serological analyses, there exists an unmet need for methods that can characterize antibodies to post-translationally modified or haptenized epitopes. Here, we adapted the antibody profiling platform PhIP-Seq to the quantitative analysis of antibody reactivity towards an enzymatically citrullinated human peptidome library. This allowed us to characterize and track the fine specificities of ACPA responses of individual RA patients over time, without bias, in high-throughput and at relatively low cost. The use of PTM PhIP-Seq is broadly applicable to studies of diverse PTMs, potentially adaptable to studies of glycosylation,^41^ carbamylation,^42^ phosphorylation,^43^ deamidation,^44^ and other modifications.^6^ PTMs of the phage displayed human peptidome may also be used to profile protease activity via the SEPARATE assay.^45,46^ PTM PhIP-Seq is therefore a powerful addition to the programmable phage display toolkit with broad utility for genome-wide proteomic analyses.

## Methods

### Human subjects

Sera from 30 individuals with RA from a cross-sectional cohort and 10 demographically matched healthy controls were studied. Patient serum was collected under a study protocol approved by the Office of Human Subjects Research Institutional Review Boards of the Johns Hopkins University School of Medicine; written consent was obtained prior to participation. Healthy control sera were obtained from volunteers who self-identified, were not pregnant, and did not have a history of cancer, autoimmune disease, or active tuberculosis/HIV/hepatitis infection. RA patients fulfilled 2010 ACR classification criteria for RA, as described.^47^ Anti-CCP antibody levels were determined by the QUANTA Lite^®^ CCP3 IgG ELISA (Inova Diagnostics, # 704535). For one RA patient, serum was available at several time points pre- and post-treatment with the chimeric anti-CD20 monoclonal antibody rituximab. At baseline (day 0), the patient was treated with methotrexate and glucocorticoids with insufficient control of arthritis symptoms. Two doses of rituximab were administered on days 22 and 35, respectively. On day 0, anti-CCP and RF normalized units were 252 and 139 respectively.

### Expression and purification of peptidylarginine deiminases

Recombinant human PAD4 and PAD2 containing N-terminal 6 x histidine tags were expressed in BL21(DE3) pLysS competent cells and purified by immobilized metal affinity chromatography with a nickel resin, as previously described.^48,49^

### Phage modification quality control

A negative control clone that lacks a displayed peptide (due to a premature stop codon, introduced as a frameshift mutation of the DNA insert) and a positive control clone that displays a 90-mer arginine-containing peptide from POU3F1 were isolated by randomly picking plaques from the human peptidome phage library. Peptide sequences were determined by Sanger sequencing. PAD modification was performed as described below. Dialysis, plaque assay, and western blotting were performed according to standard molecular biology protocols. Histone citrullination was detected using anti-citrullinated histone antibody (citrulline R2 + R8 + R17, Abcam #ab5103) 1:1,000 in TBST and visualized by chemiluminescence. Quantitative PCR assays were performed using SYBR green on a 1:1,000 dilution of product obtained from 10 cycles of PCR1.^1^

### Human peptidome T7 phage library citrullination

Citrullination was carried out in 42 mL containing buffer alone or 150 μg of either recombinant PAD4 or recombinant PAD2, 10^12^ pfu human peptidome T7 phage library (pre-dialyzed into 100 mM Tris-HCl, pH 7.5 using a Spectra/Por 7 RC 50 kDa molecular weight cut off dialysis membrane, Spectrum Labs, #132130), 5 mM CaCl_2_ and 2 mM DTT. The reactions were incubated at 37 °C for 3 hours with end-over-end rotation and quenched by dialysis against 100 mM Tris-HCl, pH 7.5 in order to remove free Ca^2+^ ions using a 10 kDa molecular weight cutoff dialysis membrane (Thermo Fisher, #88245). A second round of dialysis against TBS, pH 7.4 was performed in order to prepare the libraries for antibody binding and immunoprecipitation. Phage viability after modification was assessed by plaque assay.

### Phage ImmunoPrecipitation Sequencing (PhIP-Seq)

Screening, PCR, and peptide read count data generation was performed as previously described.^1^ Briefly, 0.2 μL of sample (~2 μg IgG) was added to the phage library (10^10^ pfu) in TBS, pH 7.4. The phage/serum mixtures were rotated overnight at 4°C after which 40 μL of a 1:1 Protein A/G coated magnetic bead mixture (Invitrogen, #10002D and #10004D) was added to each well and rotated for an additional 4 hours at 4°C to capture all IgG. The beads were then washed three times with TBS, pH 7.4 containing 0.1% NP-40 and resuspended in 20 μL of a Herculase II Fusion Polymerase (Agilent, #600679) PCR1 master mix to amplify library inserts. Sample-specific barcoding and the Illumina P5/P7 adapters were then incorporated during a subsequent PCR2 reaction. PCR2 amplicons were pooled and sequenced using an Illumina NextSeq 500 SE 1×50 protocol.

### Pairwise analysis of PTM PhIP-Seq data

We used pairwise enrichment analysis to identify peptides that were differentially reactive between treatment conditions. Robust regression of the top 1,000, by abundance, unmodified read counts (Unmod) was used to calculate the ‘expected’ modified read counts (PAD2 & PAD4). The observed modified read counts minus the expected modified read counts for each peptide was calculated to determine peptide residuals. Peptides were grouped in bins and a standard deviation was calculated between all peptides in each bin. From these binned standard deviations, a linear regression was developed and used to assign each peptide an expected standard deviation. Each peptide’s residual was normalized to its expected standard deviation, in order to calculate a ‘pairwise z-score’. Peptide reactivities were considered positive for pairwise z-scores ≥7.

### Statistical analyses

Differences among the total number of peptide enrichments between sample groups was calculated using a Wilcoxon test (**Fig. 4**). Differences among peptide read counts were calculated using a Kruskal-Wallis test with Dunn’s correction (**Fig. S4**). Differences between the total number of anti-CCP+ patient specific PAD2- and PAD4-unique reactivities were determined using an exact binomial test assuming a null probability of 0.5 for independent PAD2/4-specific peptides (**Fig. 5e**). Independent sets of peptides were determined using the maximal independent vertex set function from the R package igraph.

## Acknowledgements

This study was made possible by a National Institute of General Medical Sciences (NIGMS) grant R01 GM136724 (HBL) and funding from the Intelligence Advanced Research Projects Activity (HBL, MA). MFK was supported by the National Institute of Arthritis and Musculoskeletal and Skin Diseases (NIAMS) grant T32AR048522. We are grateful to Stephen J. Elledge (Harvard Medical School) for generously providing the human peptidome library used in this study.

## Competing Interests

HBL is a founder of Portal Bioscience, Alchemab and ImmuneID, and is an advisor to TScan Therapeutics. ED is author on a licensed patent (US patent no. 8,975,033) and licensed provisional patent (US patent no. 62/481,158) related to the use of antibodies to PAD3 and PAD2, respectively, in identifying clinically informative disease subsets in RA, has received consulting fees from Celgene and Bristol Myers Squibb, and has received research support from Pfizer, Celgene, and Bristol Myers outside of this work.

## Author Contributions

Conceptualization: GDRM, RSQ, MA, ED, HBL; Data curation: GDRM, DRM; Formal analysis: GDRM, DRM, MFK, ED, HBL; Funding acquisition: MA, HBL; Methodology: GDRM, HBL; Resources: JMM, RSQ, MFK, MA, ED, HBL; Software: DRM; Visualization: GDRM, DRM, MFK, ED, HBL; Writing - original draft: GDRM, HBL; Writing - review and editing: DRM, JMM, MFK, MA, ED, HBL.

## Supplemental Figures

**Figure S1.**
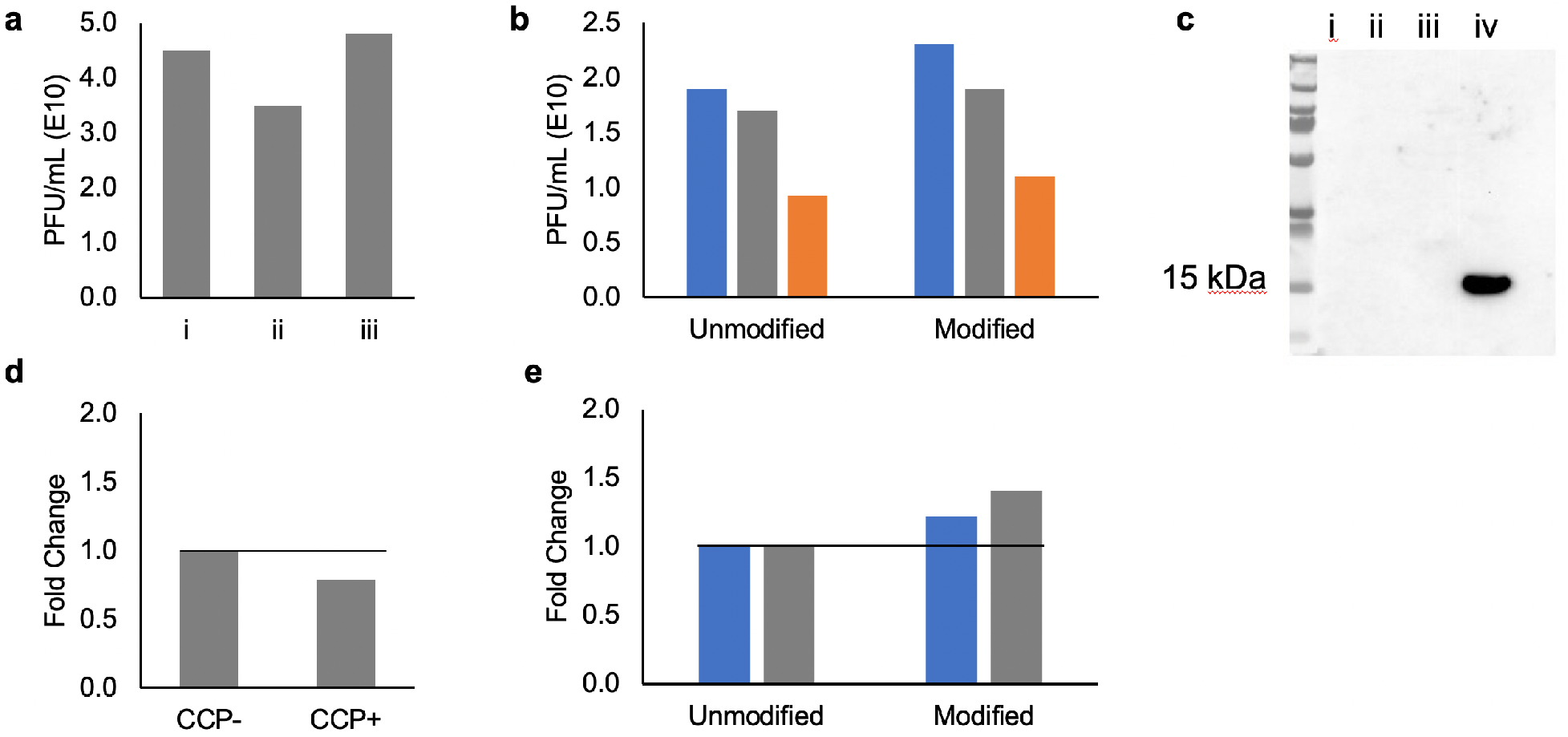
Quality control of citrulline modified human peptidome phage. **a** Phage titers pre- (i) and post- (ii, TBS; iii, PBS) dialysis are similar. **b** Phage titers pre- and post-PAD4 modification are similar for the 90-mer library (blue), no display clone (grey), non-PAD substrate clone (orange). **c** Post-PAD4 modification western blot using an anti-citrulline histone antibody was used to detect citrullination of purified histone (i, 90mer library; ii, no-display clone; iii, single clone from the 90mer library; iv, histone). **d** anti-CCP+ serum does not react with coat protein of PAD4-treated phage. Data normalized to anti-CCP- (solid line). **e** Modification does not interfere with the ability to PCR amplify library inserts from phage immunoprecipitated with anti-CCP+ (blue) or healthy control (grey) sera. PCR amplification of the phage library inserts was quantified by qPCR and the data normalized to unmodified (solid line).

**Figure S2.**
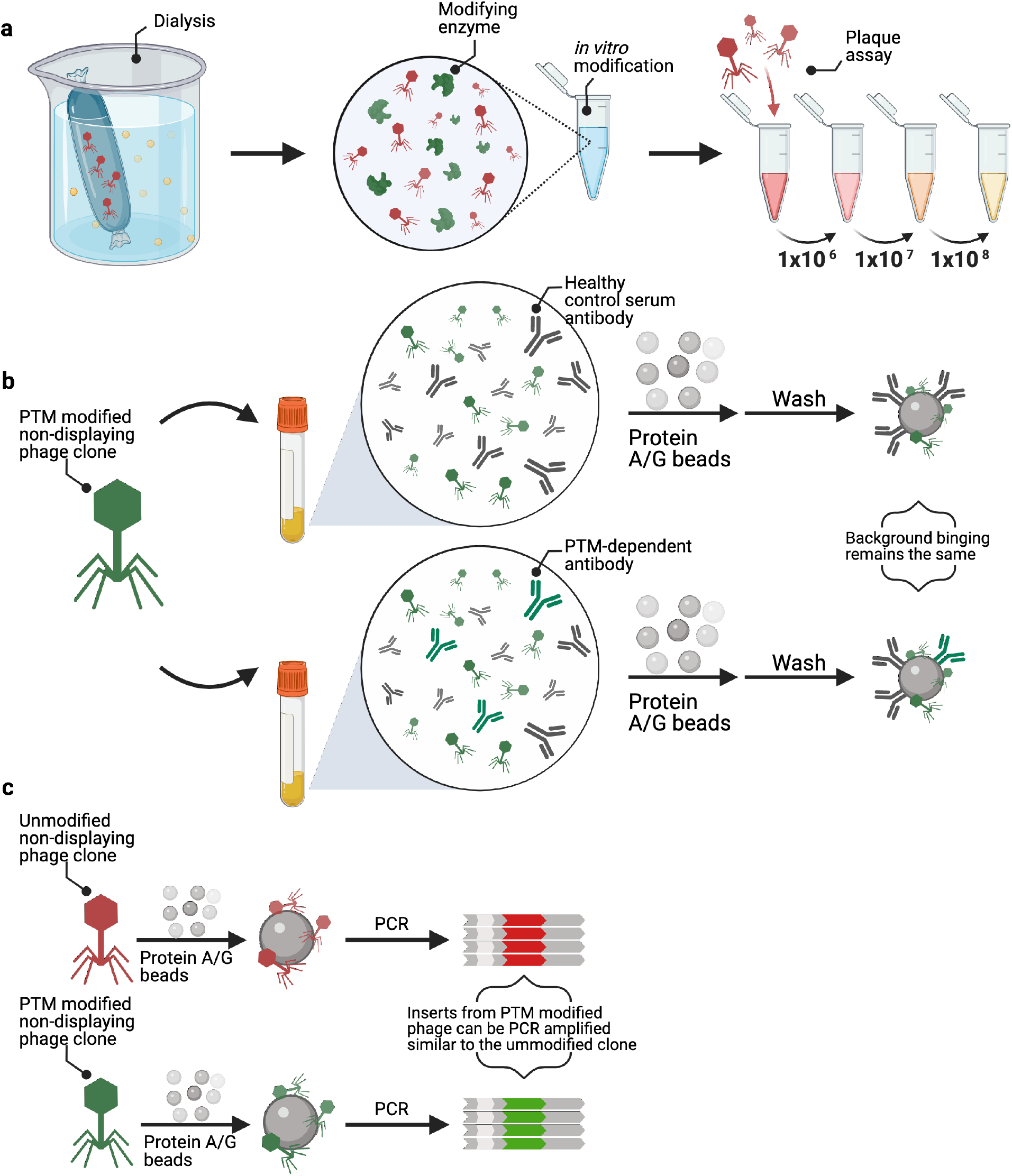
Generalizable pipeline for the quality control of PTM phage libraries. **a** Dialysis into reaction buffer and *in vitro* PTM of the phage library is followed by a plaque assay to quantify phage viability. **b** A PTM phage clone that does not display a peptide is independently screened against a healthy control and serum containing an anti-PTM antibody to ensure any generated phage coat epitopes are not recognized by anti-PTM antibodies. **c** Unmodified and PTM modified phage clones that do not display a peptide are immunoprecipitaded on protein A/G beads and the background binding is quantified to ensure modification does not interfere with the ability to PCR amplify library inserts.

**Figure S3.**
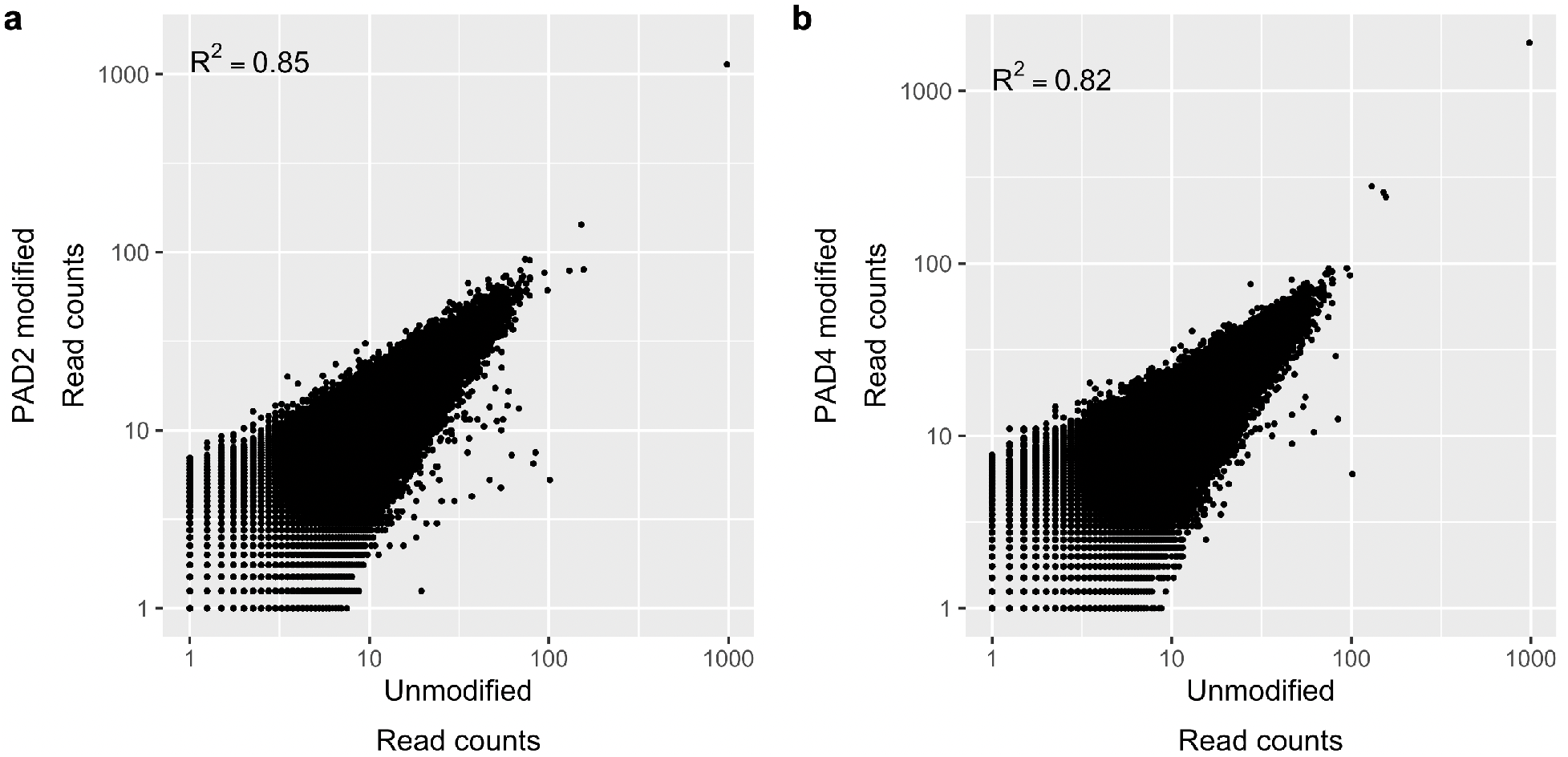
Mock immunoprecipitations in which serum was not added to the unmodified library input, compared with mock IPs using **a** PAD2 and **b** PAD4-modified libraries as input. The coefficient of determination (R^2^) was determined using linear regression.

**Figure S4.**
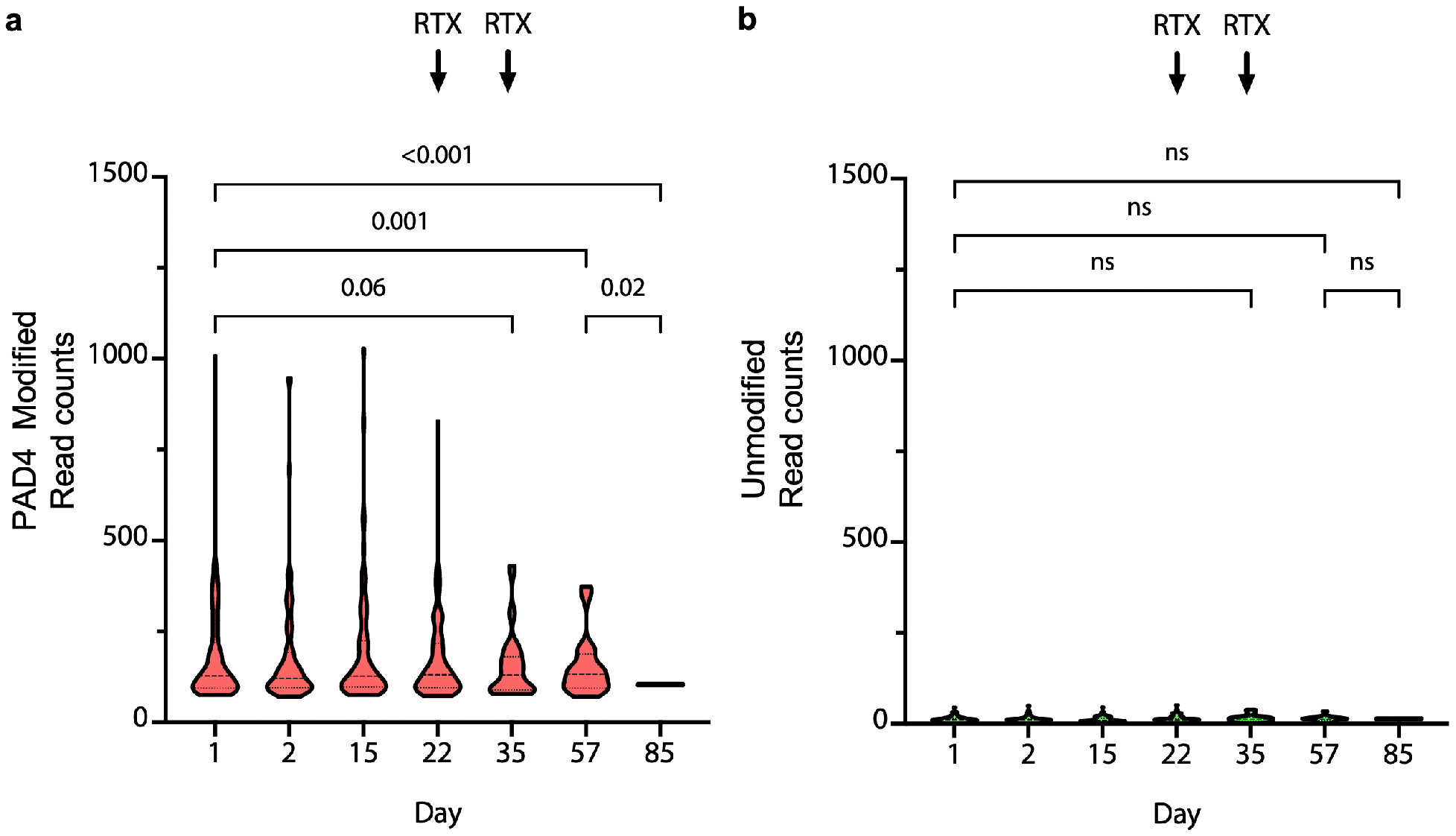
Distribution of PAD4 PhIP-Seq read counts for modified and unmodified reactive peptides from a longitudinally monitored patient with RA (same as Fig. 3). **a** The read count of each PAD4-reactive peptide declines following treatment with rituximab (RTX). **b** The read count for each analogous peptide in the unmodified library remains unchanged.

**Figure S5.**
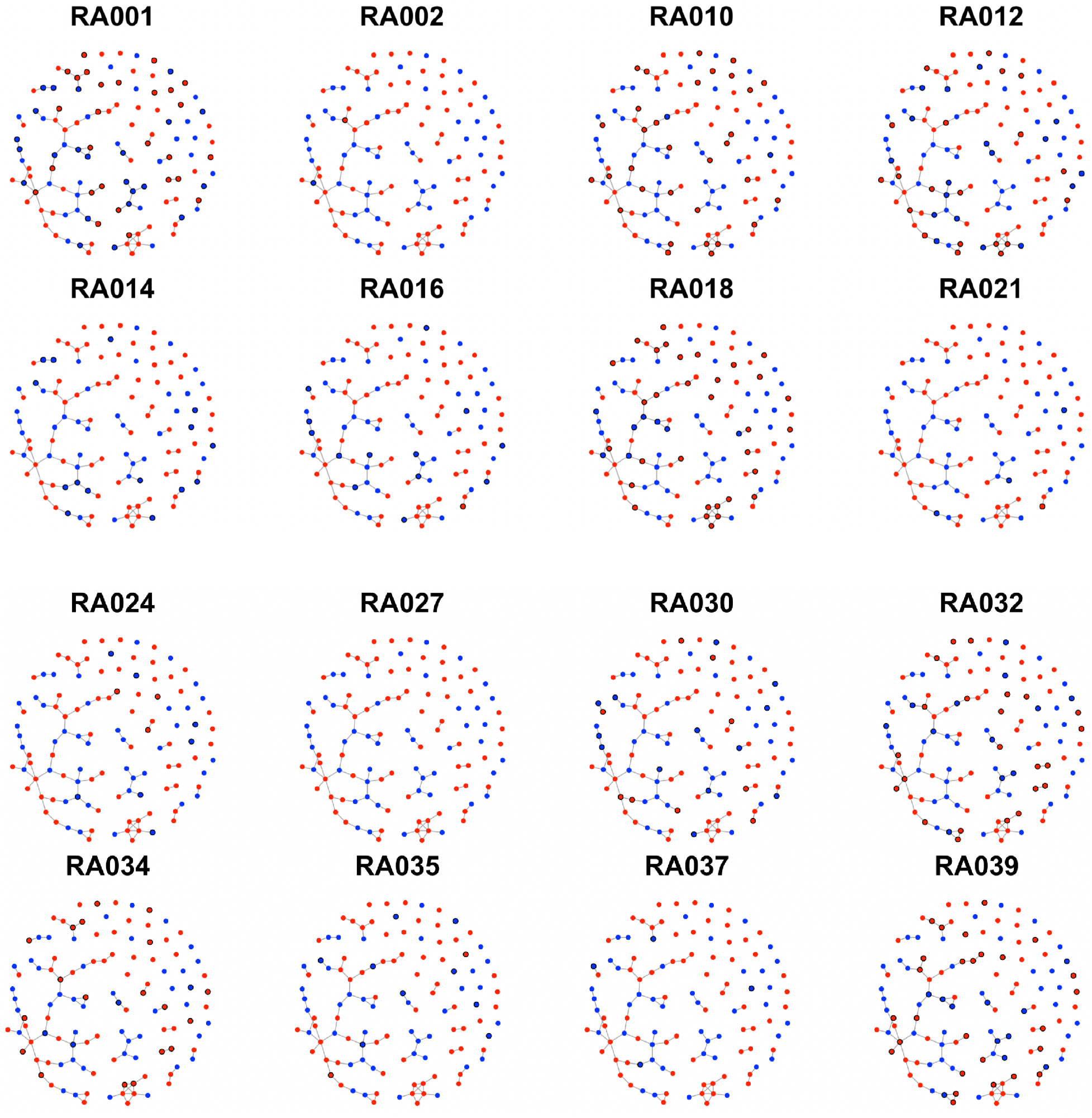
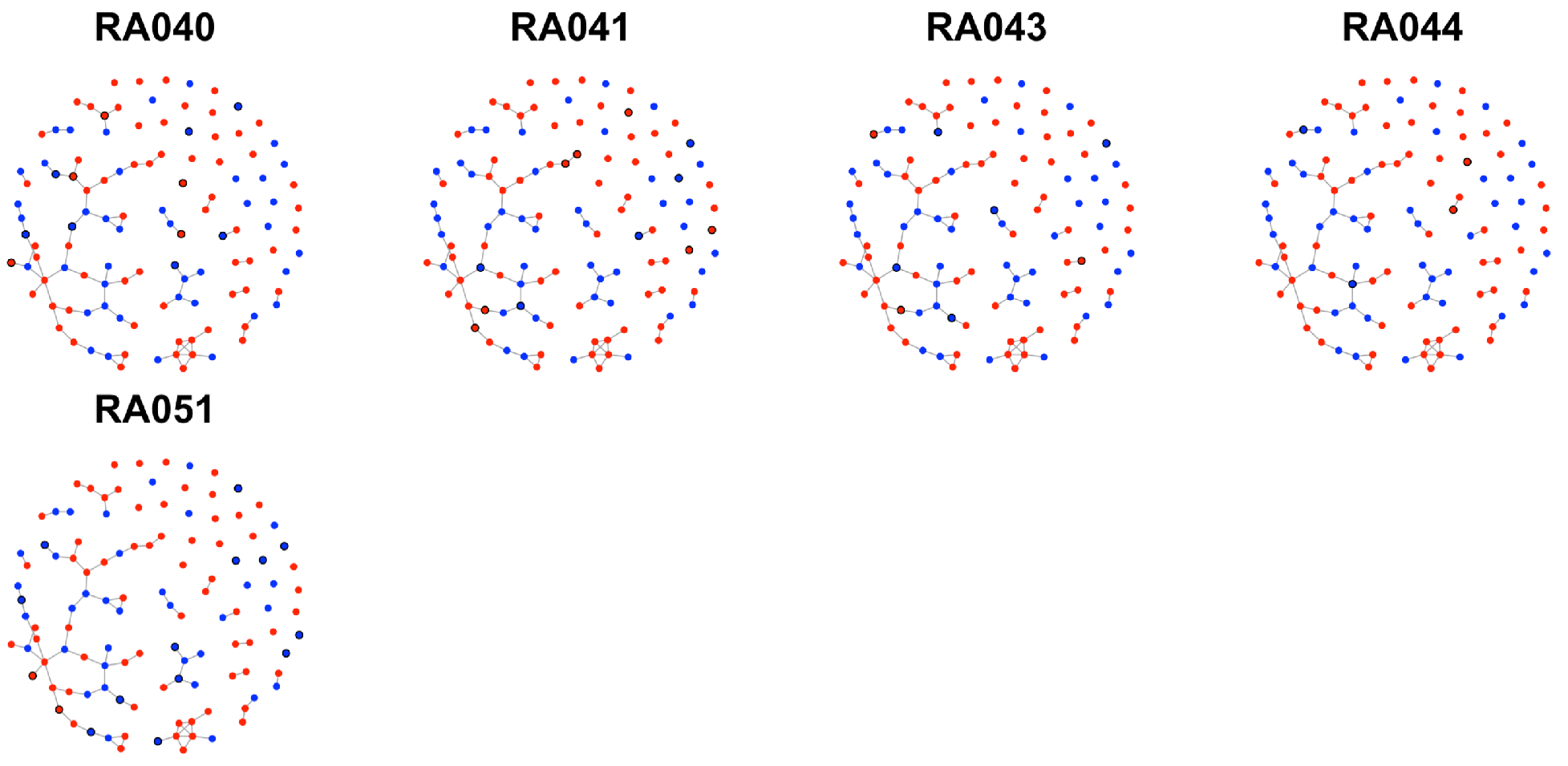
Network graph of PAD specific reactivity profiles for each anti-CCP+ individual. PAD2 specific reactivities are shown in blue. PAD4 specific reactivities are shown in orange. Each unique reactive peptide identified in each anti-CCP+ individual is circled in black.

**Table S1.** Concordance of PAD PhIP-Seq versus CCP status. A cutoff of ≥ 15 reactive peptides was used to determine seropositivity by PAD PhIP-Seq. RA: diseased sample; HC: healthy control. *Average of independent replicas: RA44: PAD2=88, 57; PAD4=61, 56; **Sjogren’s: PAD2=6, 7; PAD4=4, 4.

| Sample | Number of reactive peptides |  | CCP status | CCP units |
| --- | --- | --- | --- | --- |
|  | PAD2 PhIP-Seq | PAD4 PhIP-Seq |  |  |
| RA01 | 876 | 783 | (+) | 161 |
| RA02 | 28 | 34 | (+) | 155 |
| RA10 | 771 | 414 | (+) | 155 |
| RA12 | 313 | 349 | (+) | 163 |
| RA14 | 62 | 183 | (+) | 94 |
| RA16 | 79 | 145 | (+) | 152 |
| RA18 | 3640 | 2193 | (+) | 156 |
| RA21 | 69 | 91 | (+) | 161 |
| RA24 | 156 | 304 | (+) | 98 |
| RA27 | 4 | 1 | (+) | 163 |
| RA30 | 327 | 356 | (+) | 163 |
| RA32 | 415 | 467 | (+) | 161 |
| RA34 | 571 | 405 | (+) | 138 |
| RA35 | 104 | 162 | (+) | 162 |
| RA37 | 12 | 43 | (+) | 122 |
| RA39 | 812 | 584 | (+) | 152 |
| RA40 | 247 | 376 | (+) | 163 |
| RA41 | 418 | 485 | (+) | 162 |
| RA43 | 141 | 122 | (+) | 156 |
| RA44 | 73* | 59* | (+) | 158 |
| RA51 | 294 | 306 | (+) | NA |
| RA03 | 14 | 4 | (-) | 6 |
| RA04 | 8 | 2 | (-) | 6 |
| RA09 | 4 | 4 | (-) | 7 |
| RA13 | 2 | 2 | (-) | 6 |
| RA15 | 5 | 1 | (-) | 18 |
| RA17 | 7 | 8 | (-) | 14 |
| RA19 | 2 | 4 | (-) | 6 |
| RA20 | 13 | 4 | (-) | 7 |
| RA25 | 14 | 0 | (-) | 9 |
| RA26 | 6 | 8 | (-) | 8 |
| HC35 | 13 | 10 | NA | NA |
| HC36 | 9 | 5 | NA | NA |
| HC05 | 3 | 1 | NA | NA |
| HC11 | 10 | 2 | NA | NA |
| HC39 | 13 | 10 | NA | NA |
| HC48 | 11 | 1 | NA | NA |
| HC54 | 4 | 0 | NA | NA |
| HC56 | 5 | 0 | NA | NA |
| HC58 | 7 | 7 | NA | NA |
| HC52 | 6 | 2 | NA | NA |
| Sjogren's | 7** | 4** | NA | NA |

**Table S2.** PAD2- and PAD4-preferred reactivities. Total peptides is the sum of all PAD2 and PAD4 reactivities identified by PAD PhIP-Seq. PAD2- and PAD4-unqiue is the total number of PAD specific peptides identified for each anti-CCP+ individual. PAD2- and PAD4-test is the number of independent peptides used in the binomial test.

| Sample | Total peptides | PAD2 unique | PAD4 unique | PAD2 test | PAD4 test | p-value |
| --- | --- | --- | --- | --- | --- | --- |
| RA001 | 1659 | 27 | 23 | 24 | 19 | 5.42E-01 |
| RA002 | 62 | 1 | 1 | 1 | 1 | 1.00E+00 |
| RA010 | 1185 | 32 | 2 | 26 | 2 | 3.00E-06 |
| RA012 | 662 | 20 | 18 | 17 | 16 | 1.00E+00 |
| RA014 | 245 | 0 | 15 | 0 | 13 | 2.44E-04 |
| RA016 | 224 | 1 | 13 | 1 | 12 | 3.42E-03 |
| RA018 | 5833 | 32 | 8 | 27 | 7 | 8.21E-04 |
| RA021 | 160 | 1 | 3 | 1 | 3 | 6.25E-01 |
| RA024 | 460 | 3 | 7 | 3 | 7 | 3.44E-01 |
| RA027 | 5 | 0 | 0 | 0 | 0 | 1.00E+00 |
| RA030 | 683 | 9 | 13 | 8 | 12 | 5.03E-01 |
| RA032 | 882 | 23 | 12 | 18 | 11 | 2.65E-01 |
| RA034 | 976 | 19 | 4 | 16 | 4 | 1.18E-02 |
| RA035 | 266 | 2 | 9 | 2 | 9 | 6.54E-02 |
| RA037 | 55 | 0 | 4 | 0 | 4 | 1.25E-01 |
| RA039 | 1396 | 27 | 11 | 22 | 9 | 2.94E-02 |
| RA040 | 623 | 5 | 7 | 5 | 7 | 7.74E-01 |
| RA041 | 903 | 7 | 5 | 6 | 5 | 1.00E+00 |
| RA043 | 263 | 3 | 5 | 3 | 5 | 7.27E-01 |
| RA044 | 149 | 2 | 2 | 2 | 2 | 1.00E+00 |
| RA051 | 600 | 2 | 13 | 2 | 13 | 7.39E-03 |

